# Altered renal vascular patterning reduces ischemic kidney injury and limits vascular loss associated with aging

**DOI:** 10.1101/2024.10.29.620969

**Authors:** Sarah R. McLarnon, Samuel E. Honeycutt, Pierre-Emmanuel Y. N’Guetta, Yubin Xiong, Xinwei Li, Koki Abe, Hiroki Kitai, Tomokazu Souma, Lori L. O’Brien

## Abstract

The kidney vasculature has a complex arrangement, which runs in both series and parallel to perfuse the renal tissue and appropriately filter plasma. Recent studies have demonstrated that the development of this vascular pattern is dependent on netrin-1 secreted by renal stromal progenitors. Mice lacking netrin-1 develop an arterial tree with stochastic branching, particularly of the large interlobar vessels. The current study investigated whether abnormalities in renal vascular pattern altered kidney function or response to injury. To examine this, we analyzed kidney function at baseline as well as in response to recovery from a model of bilateral ischemic injury and measured vascular dynamics in aged mice. We found no differences in kidney function or morphology at baseline between mice with an abnormal arterial pattern compared to control. Interestingly, male and female mutant mice with stochastic vascular patterning showed a reduction in tubular injury in response to ischemia. Similarly, mutant mice also had a preservation of perfused vasculature with aging compared to a reduction in the control group. These results suggest that guided and organized patterning of the renal vasculature may not be required for normal kidney function; thus, modulating renal vascular patterning may represent an effective therapeutic strategy. Understanding how patterning and maturation of the arterial tree affects physiology and response to injury or aging has important implications for enhancing kidney regeneration and tissue engineering strategies.

## Introduction

The kidney vasculature is structurally complex to appropriately filter waste products from the blood and maintain water and electrolyte homeostasis^1–4^. In the mouse kidney, the main renal artery is derived via angiogenic growth from the abdominal aorta, forming a ring around the base of the developing ureteric bud around embryonic day 11 (E11)^5^. Through a series of growth and branching, the endothelium of the arterial tree forms a predictable pattern of vessels in the developing kidney by E13.5^6^. This pattern consists of a single renal artery which branches in series after entering the hilum into interlobar, arcuate and interlobular arteries^3,4^. The afferent and efferent arteries then supply blood to and remove blood from the glomeruli^4^. A dense capillary network perfuses the cortex (peritubular capillaries) and medulla (plexus capillaries)^3,4^. The venous system tracks alongside the arterial tree to remove blood from the kidney^4^. The maturation of the vascular tree throughout nephrogenesis is dependent on the development of a variety of mural cells which closely interact with the vascular endothelium^6,7^. Vascular smooth muscle cells (VSMCs) cover the larger arterial branches, while pericytes stabilize the capillary networks of the cortex and medulla^8–10^. The glomerular capillaries mature through interactions with adjacent podocyte and mesangial cells precursors^10^. While recent studies have begun to elucidate the molecular signaling required to direct this precise patterning, the effect of renal vascular patterning on kidney function and physiology remains unknown.

The vascular network in the kidney is critical to its function, and disruptions of these intricate networks can lead to the onset or progression of kidney diseases^11,12^. Loss of vascular density, or capillary rarefaction, is associated with several kidney-related diseases including hypertension, diabetic kidney disease, and acute kidney injury (AKI)^13–16^. Vascular rarefaction or vascular dysfunction has been shown to promote regional ischemia, impair renal hemodynamics and promote the development of interstitial fibrosis and chronic kidney disease (CKD)^11–13^. Preventing capillary drop-out or promoting regeneration are promising therapeutic strategies to prevent or slow progression of functional decline in the kidney^11^. As such, recent efforts have been made to specifically target the renal vasculature in CKD including inflammation, endothelial-to-epithelial crosstalk, angiogenic factors, mural cell coverage as well as others^11,17–19^. While vascular targeting strategies show promising results in rodent models, limitations remain for the implementation of these therapies in the clinical setting. In addition to these *in vivo* models, bioengineering of vascularized kidneys has also been a primary strategy in regeneration efforts. Similarly, efforts to appropriately vascularize these tissues have remained incomplete^20^. A critical limitation to both vascular targeting strategies and bioengineering kidneys is our lack of knowledge regarding whether correct, guided vascular patterning is crucial for kidney function, injury, and repair.

Our understanding of how the kidney vascular pattern affects kidney function has been limited, in part, due to our lack of animal models reflecting disrupted patterning. We recently reported that loss of netrin-1 (*Ntn1*) from Foxd1+ stromal progenitors in the kidney results in abnormal patterning of the large caliber vessels, such as the interlobar vessels, during kidney development without affecting the overall vascular density^8^. Netrin-1 (*Ntn1*) is a known guidance cue for axonal growth that is highly expressed by the kidney stromal progenitors during nephrogenesis^8,9,21,22^. In addition, netrin-1 has been shown to regulate vessel growth, branching, and organizational patterning by generating chemo-attractive or repulsive cues through receptors CD146 and Unc5b, respectively^23–25^. Our previous studies revealed a similar role for netrin-1 in vascular guidance in the developing kidney, where secretion of netrin-1 by the stromal progenitor cells was required for proper vascular patterning and vascular maturation^8^. Similar to our findings, Luo *et al.* have also reported dysregulation in arterial branching and delayed VSMC coverage following deletion of *Ntn1* from the stromal progenitor cells^9^. Taken together, these studies are the first to highlight a role for stromal progenitor-derived netrin-1 (*Ntn1*) to direct renal vascular patterning and maturity during kidney development. This new model is ideal to dissect the impacts of large vascular patterning on kidney function.

It has previously been reported that the large renal vasculature (interlobar/arcuate vessels) may protect the tubular epithelial cells from ischemic injury by altering reperfusion hemodynamics^3^. Therefore, in the current study, we investigated the role of arterial patterning on kidney function and injury. We utilized *Foxd1^GC/+^;Ntn1^fl/fl^*mutant mice in which arterial patterning is disrupted during development and persists into adulthood^8,9^. Ischemia-reperfusion injury (IRI) was performed in male and female mice to assess their response to ischemic injury. Further, we examined how vascular patterning contributes to age-related losses in vascular density. Understanding how vascular patterning and maturity regulate kidney physiology is important to identify new therapeutic targets to prevent or limit vascular-associated damage following ischemia or during aging.

## Results

### Abnormal renal vascular patterning does not alter baseline kidney function or histology in adult mice

To investigate the role of abnormal renal vascular patterning, occurring during nephrogenesis, on kidney physiology, we first assessed baseline parameters of kidney function in 4-month-old adult wild-type (*Ntn1^fl/fl^* or *Ntn1^fl/+^*), heterozygous (*Foxd1^GC/+^;Ntn1^fl/+^*) and mutant (*Foxd1^GC/+^;Ntn1^fl/fl^*) male and female mice. We found that despite kidneys from mutant mice having stochastic patterning of the arterial tree, there was no evidence of altered kidney function or kidney injury at baseline. Proteinuria (**Figure 1A**) and plasma creatinine (pCr) (**Figure 1B**) were within normal physiological range (proteinuria <2mg/day and pCr <0.25mg/dL) regardless of sex or genotype. Baseline glomerular filtration rate (GFR, µl/min) measured by creatinine clearance showed no significant differences between genotypes (**Figure 1C**). Urine output (ml, 24-hours) matched water intake (ml, 24-hour) in male mice regardless of genotype (**Figure 1D-E**, gray bars). In female mice, urine volume (ml) was less than water intake for all genotypes (**Figure 1D-E**, white bars). In addition, female mice were smaller than male mice, regardless of genotype (**Figure 1F**). Heterozygous (*Foxd1^GC/+^;Ntn1^fl/+^*) and mutant female mice (*Foxd1^GC/+^;Ntn1^fl/fl^*) had reduced body weight (g) compared to the wild-type control (*Ntn1^fl/fl^* or *Ntn1^fl/+^*), however this was not observed in male mice (**Figure 1F**).

**Figure 1.**
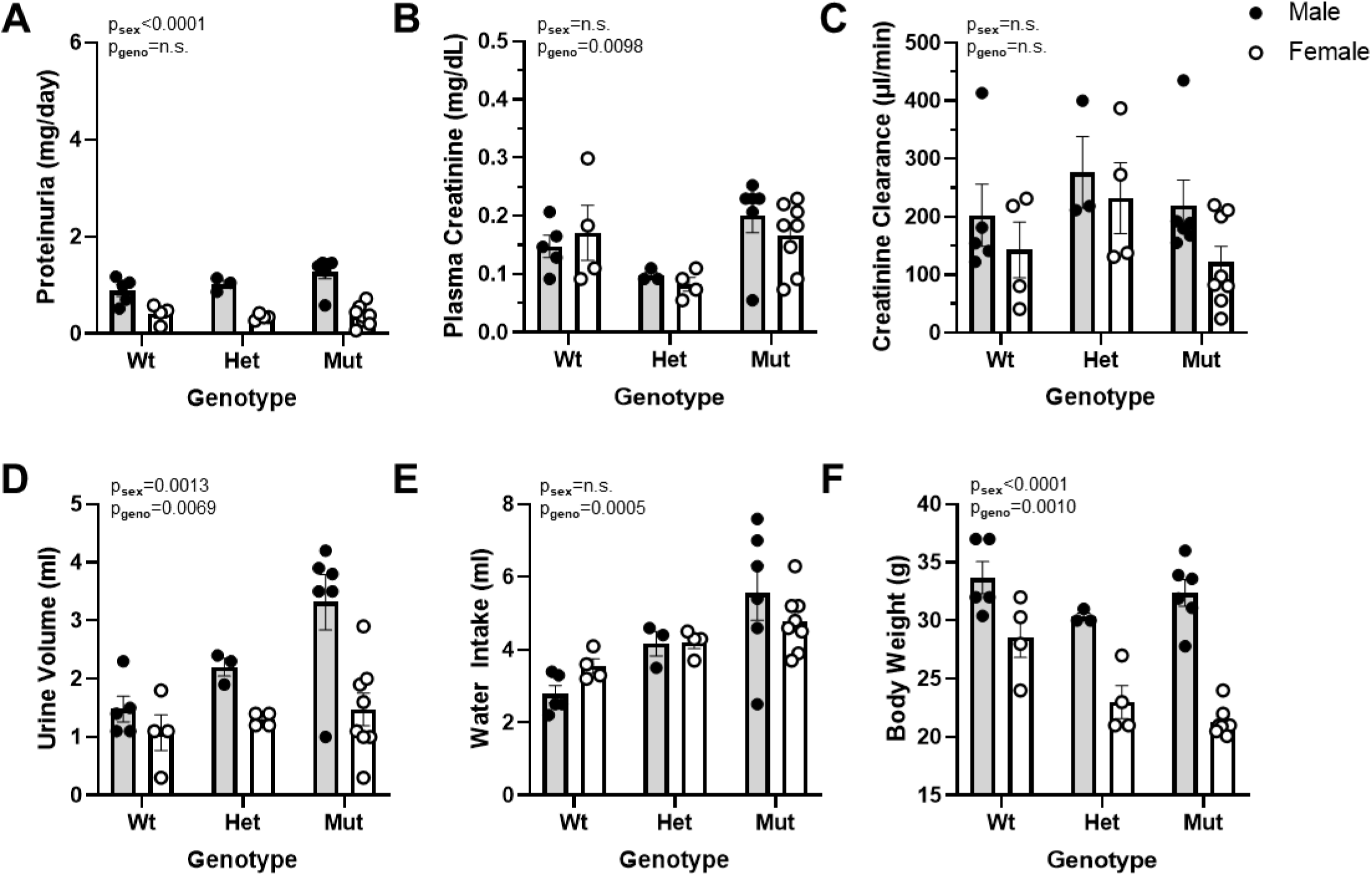
Netrin-1 mutants with abnormal patterning of the renal vasculature exhibit normal baseline kidney function. Proteinuria (mg/day) (**A**), plasma creatinine (mg/dL) (**B**), creatinine clearance (µl/min) (**C**), urine volume (ml) (**D**), water intake (ml) (**E**), and body weight (g) (**F**) in wild-type (Wt, *Ntn1^fl/fl^* or *Ntn1^fl/+^*), heterozygous (Het, *Foxd1^GC/+^;Ntn1^fl/+^*), and mutant (Mut, *Foxd1^GC/+^;Ntn1^fl/fl^*) mice (∼4 months old). Male mice are represented in gray bars and female mice are represented in white bars. Metabolic cage parameters were collected over 24-hours following an overnight acclimation period. Values are mean±SEM, n=3-8 mice per group. Two-way ANOVA, statistical significance p<0.05.

To examine morphology at baseline, kidney sections were stained with Periodic-acid Schiff (PAS) and analyzed for abnormalities in the cortex, corticomedullary junction (CMJ), outer (OM) and inner (IM) medulla. The tubules, vasculature and interstitial space appeared normal and showed no evidence of physiological perturbations for any of the groups (**Figure 2**). There were also no signs of kidney injury as no tubular dilation, tubular necrosis, loss of brush boarder, protein cast formation or epithelial cell swelling was observed (**Figure 2**). In addition to kidney morphology, we also examined immune cells and found no differences in baseline circulating immune cells (**Supplementary Figure 2A**) or F4/80-positive macrophages in the kidney (**Supplementary Figure 2B)**. Red blood cell counts were also no different between wild-type control and mutant mice (**Supplementary Figure 3**).

**Figure 2.**
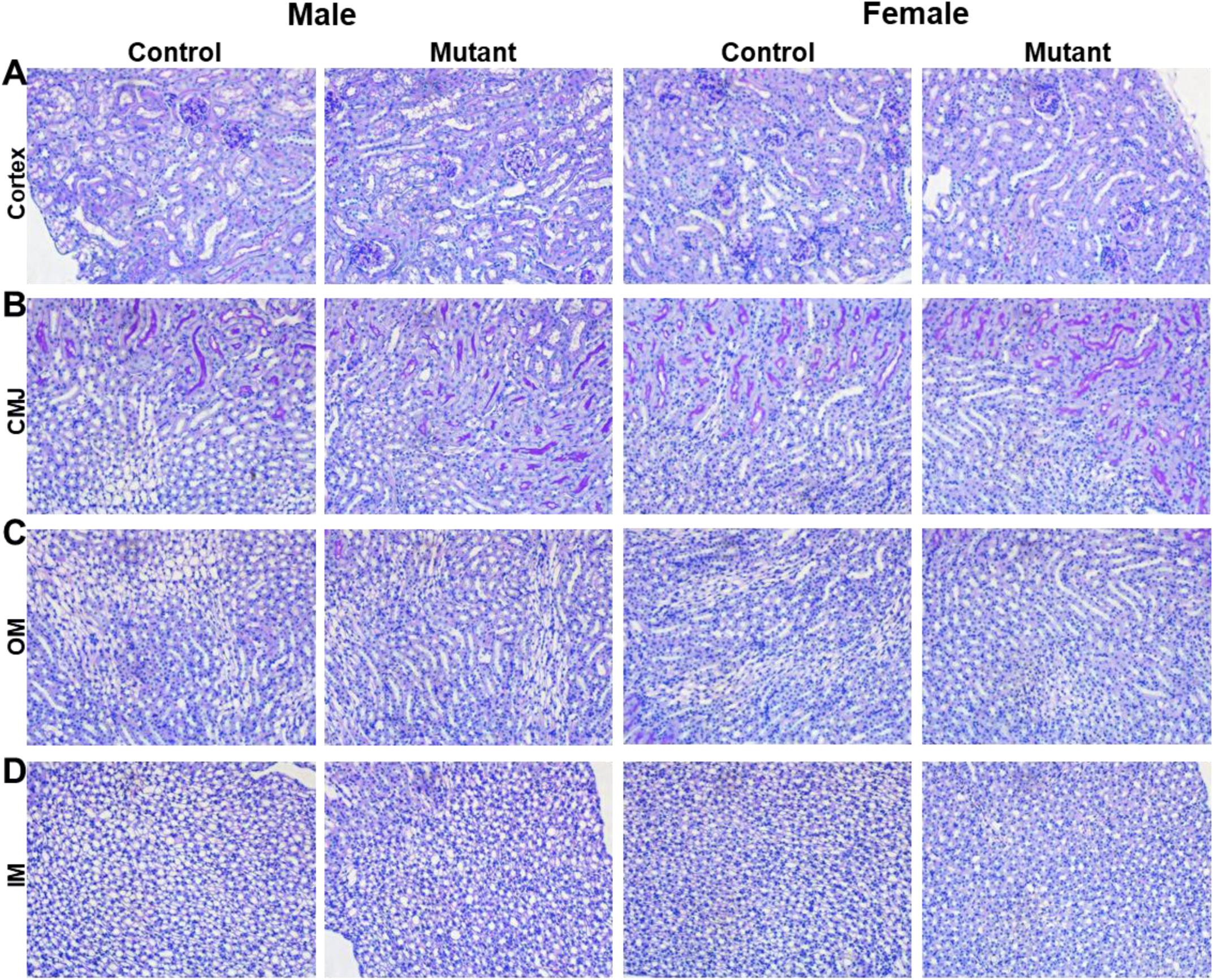
Netrin-1 mutants with abnormal patterning of the renal vasculature exhibit normal baseline kidney morphology. Representative images of baseline kidney histology from the cortex (**A**), corticomedullary junction (CMJ) (**B**), outer medulla (OM) (**C**), and inner medulla (IM) (**D**) in control (*Ntn1^fl/fl^* or *Foxd1^GC/+^;Ntn1^fl/+^*) and mutant (*Foxd1^GC/+^;Ntn1^fl/fl^*) adult mice (∼4 months old). Male mice are shown in columns 1 and 2. Female mice are shown in columns 3 and 4. Kidney sections were stained with Periodic Acid Schiff (PAS) and assessed for abnormal morphology including, tubular dilation, loss of brush boarder in the proximal tubules, tubular necrosis, protein casts, glomerular damage, and fibrosis. N=3-8 mice per group. Kidneys are from mice used for kidney function analysis in figure 1.

### Tubular injury following renal ischemia was attenuated in mice with abnormal vascular patterning

As no significant differences were observed in baseline renal function or morphology between genotypes, we next examined the renal response to ischemic injury. We performed a warm, bilateral ischemia by clamping the renal artery and vein for 26-minutes and assessed kidney injury following 24-hours of recovery. Post-IRI, the kidney to body weight ratio, which may indicate renal edema, was decreased in male, but not female mutant mice (**Supplemental Figure 3**). Tubular injury was semi-quantitatively scored in PAS-stained kidney sections on a scale of 0-5 as a percent of tubules in the cortex and outer medulla with necrosis, loss of brush borders, tubular dilatation, cast formation, tubular epithelial swelling, and vacuolar degeneration, as previously reported^26^. Interestingly, tubular injury was reduced in both male and female mutant mice compared to their respective controls despite the kidneys from these mice having an abnormal arterial pattern (p_GENO_=0.0092, two-way ANOVA) (**Figure 3A**). While all genotypes had an average injury area greater than 30% following IRI, wild-type male mice showed the greatest amount of injury (average injury score ∼75% area) (**Figure 3A&D-E**). Tubular dilation and tubular protein casts were observed in all groups, however significant tubular epithelial cell detachment/cell sloughing in the outer medulla (more severe injury) was most prominent in wild-type males (**Figure 3D-E**, asterisks and arrow). As expected, female mice had less injury than male mice regardless of genotype, and this injury was primarily localized to the outer medulla versus the cortex (p_SEX_=0.0122, two-way ANOVA) (**Figure 3A&D-E**). In agreement with these data, pCr (**Figure 3B**) and blood urea nitrogen (BUN) (**Figure 3C**) were reduced in male mutant mice compared to male control (pCr 0.46±0.16 vs 0.66±0.11 and BUN 65.61±22.54 vs 101.97±14.36, respectively), however this did not reach statistical significance.

**Figure 3.**
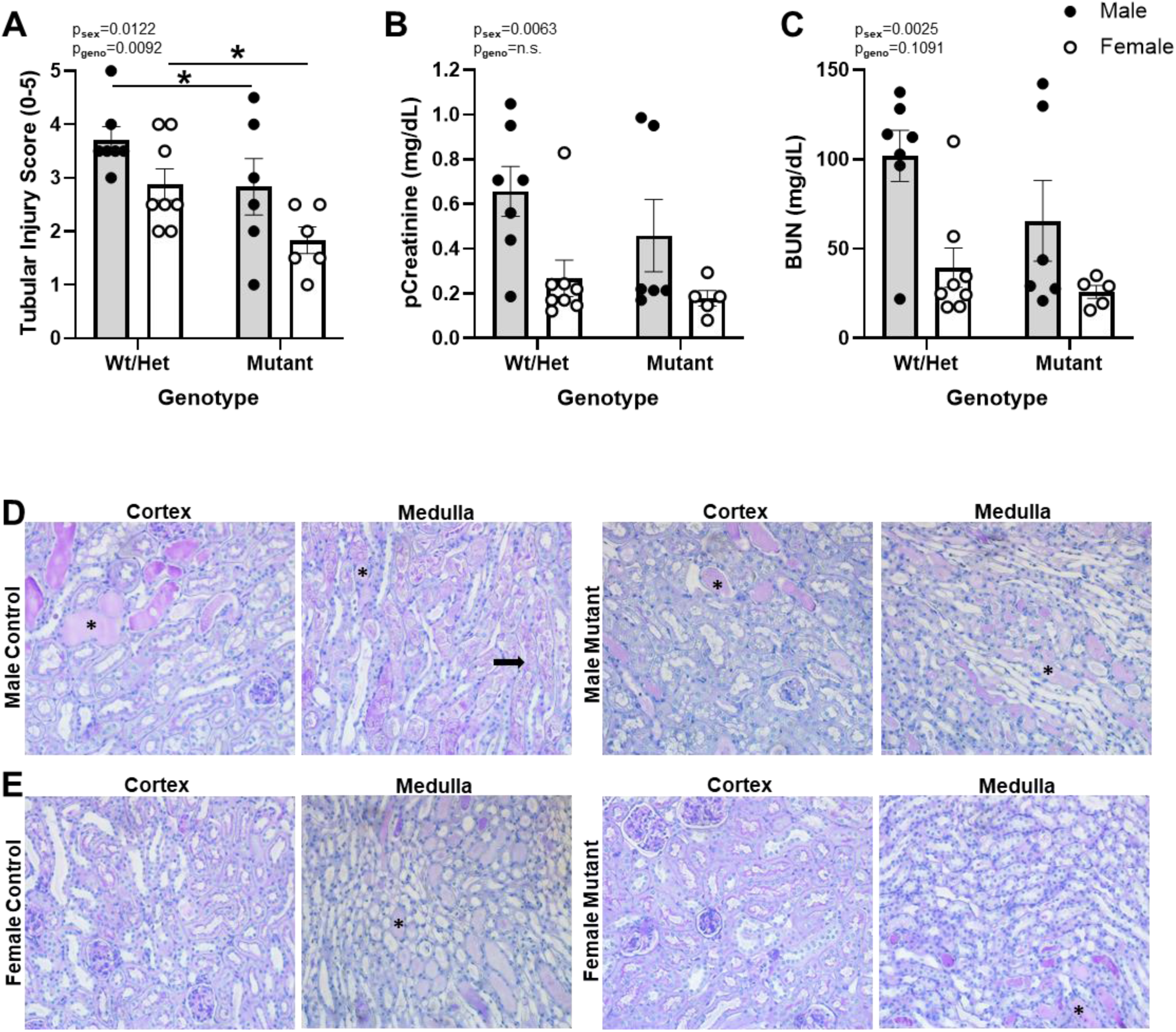
Netrin-1 mutants with abnormal patterning of the renal vasculature show a reduction in kidney injury following ischemia. Tubular injury scores (**A**) following a 26-minute bilateral ischemia and one-day of recovery in control (*Ntn1^fl/fl^* or *Foxd1^GC/+^;Ntn1^fl/+^*) and mutant (*Foxd1^GC/+^;Ntn1^fl/fl^*) adult mice (∼4 months old). Tubular injury was scored (0-5) in kidney histology sections stained with Periodic Acid Schiff (PAS). Renal damage including dead tubules, loss of brush borders, tubule dilatation, cast formation, tubular epithelial swelling, and vacuolar degeneration was semi quantitatively scored on a scale of 0-5; a score of 0 representing no injury up to a score of 5 representing >90% of kidney tissue was injured. Plasma creatinine (pCr, mg/dL) (**B**) and blood urea nitrogen (BUN, mg/dL) (**C**) were measured one-day following a 26-minute bilateral ischemia. Wild-type (WT) and heterozygous (het) mice were combined into a control group as no differences were observed. Male mice are represented in gray bars and female mice are represented in white bars. Representative histology images from PAS-stained kidney sections from the renal cortex and medulla in male (**D**) and female (**E**) control and mutant mice. Asterisks denote tubular protein casts, and the black arrow denotes tubular cell death. Values are mean±SEM, n=6-8 mice per group. Creatinine and BUN was not measured for one female mutant as not enough plasma was collected at euthanasia to run the assays. Two-way ANOVA, statistical significance p<0.05 (*).

To further investigate the reduction of injury observed in mutant mice, we utilized immunofluorescent staining of kidney injury molecule-1 (KIM-1) to assess injury to the proximal tubules (LRP2, marker of proximal tubules) in kidney sections post-IRI. KIM-1 localized to the tubules of both the cortex and outer medulla of control male mice following ischemia (**Figure 4A**). In contrast, KIM-1 staining was predominately localized to the tubules of the outer medulla and mostly absent in the cortex in mutant male mice and in both control and mutant female mice (**Figure 4B-D**).

**Figure 4.**
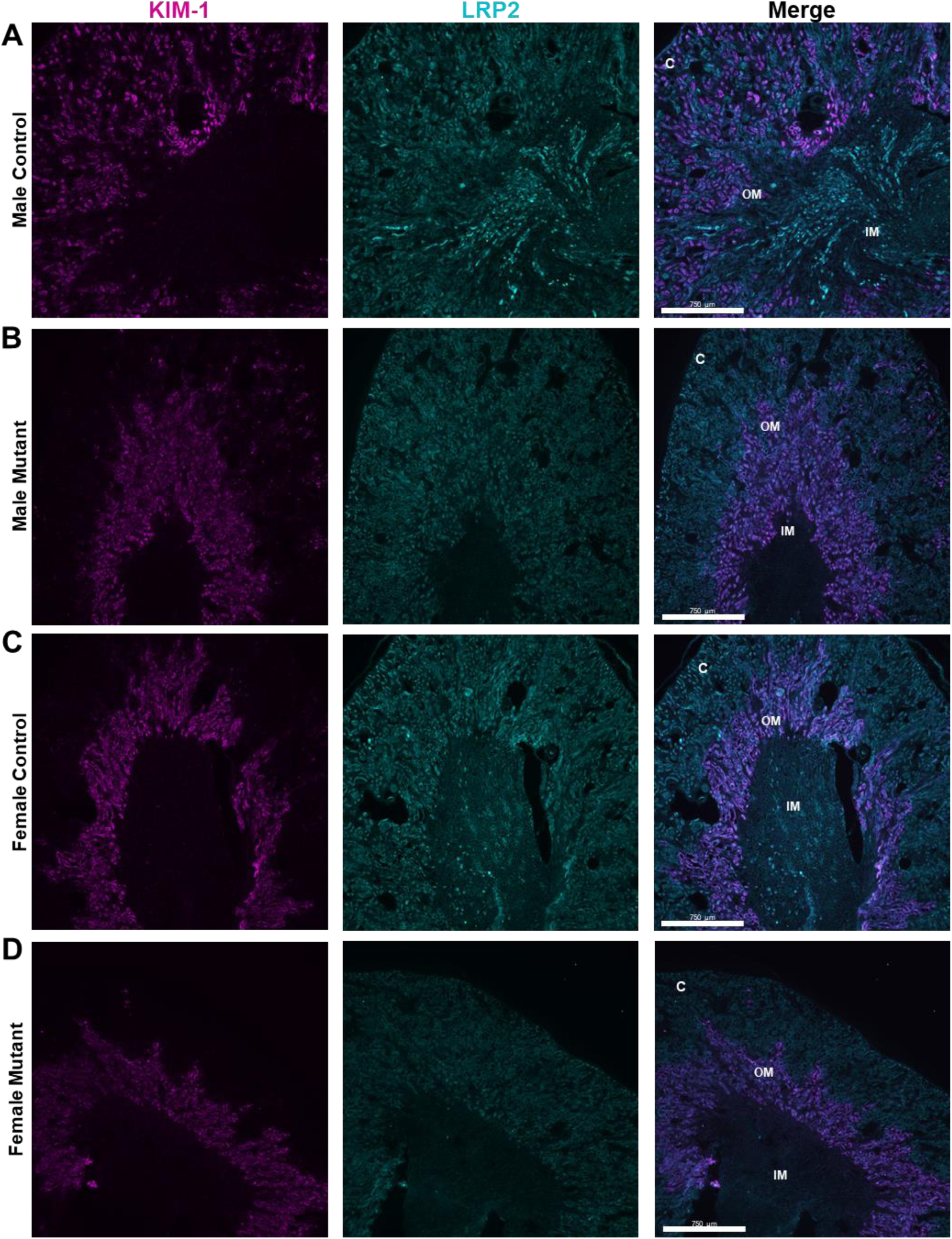
Netrin-1 mutants with abnormal patterning of the renal vasculature show attenuated injury to the proximal tubules. Representative images of immunofluorescent staining of kidney injury marker-1 (KIM-1, magenta) and megalin (LRP2, cyan) one-day following a 26-minute bilateral ischemia in control (*Ntn1^fl/fl^* or *Foxd1^GC/+^;Ntn1^fl/+^*) and mutant (*Foxd1^GC/+^;Ntn1^fl/fl^*) male (**A-B**) and female (**C-D**) mice (∼4 months old). For localization, the renal cortex (C), outer (OM), and inner (IM) medulla are labeled on the merged image. N=6-8 mice per group, kidneys are from mice used for injury analysis in figure 3. Scale bar = 750 µm. Images were captured with a Leica DMi8 inverted widefield fluorescent microscope with Leica DFC9000 GT camera and LASX software. Images were taken at 5x magnification and acquisition settings were the same for all samples. Note, KIM-1 is localized to predominately the OM except for kidneys from male control mice where staining localized to tubules in both the cortex and OM.

### Perfused renal vasculature was preserved in aged mice with abnormal vascular patterning

Our findings indicate that mutant mice with abnormal patterning of the renal vasculature have reduced tubular injury and improved kidney function following ischemia. As loss of renal vascular density following ischemia and in normal aging has been shown to contribute to the development of chronic kidney disease, we next wanted to determine whether renal vascular patterning played a role in age-related vascular impairment. To examine this, we perfused the renal vasculature with Evans blue dye in 2 and 7-month-old mice to fluorescently label the arterial tree^27^. We then cleared the kidney tissue and performed whole mount imaging using light-sheet microscopy to assess vascular parameters of branching and total area^27^. In control mice, we observed an age-related decrease in number of vascular branches (control mice: 2 vs 7-month, p=0.0043) and total perfused vasculature (control mice: 2 vs 7-month, p=0.0004) (**Figure 5A&B**, white bars). Similar to the protective effect observed following ischemia, neither the number of branches nor the total perfused vasculature significantly decreased with age in mutant mice (**Figure 5A&B**, gray bars). Such that, 7-month mutant mice had a greater area of perfused vasculature compared to age-matched control mice (approximately 70 vs 50 µM (x10^6^), respectively) (p=0.0152) (**Figure 5A&B**). Importantly, glomerular number, plasma creatine (<0.5 mg/dL) and BUN (<25 mg/dL) were not different between groups in adult mice, suggesting that an age-related decline in kidney function was not responsible for the differences in vascular metrics (**Figure 5C&D**). Circulating immune cells (**Supplementary Figure 2A**) and red blood cell counts (**Supplementary Figure 3**) also showed no differences between genotypes with aging. Taken together, these data suggest that despite mutant mice developing stochastic arterial branches in the kidney, these mice are protected from both ischemic injury and age-related loss of vascular density.

**Figure 5.**
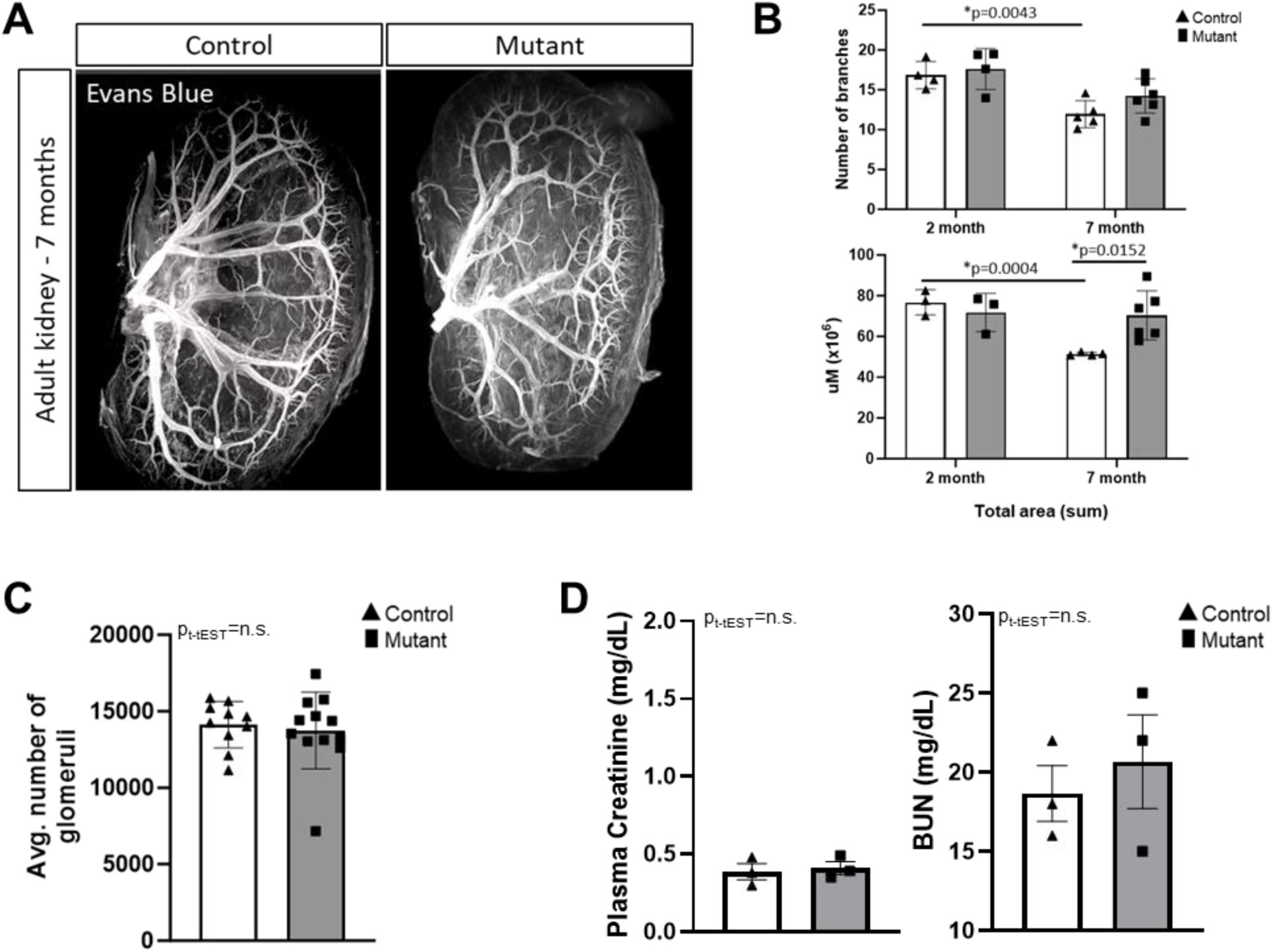
Netrin-1 mutants with abnormal patterning of the renal vasculature exhibit improved preservation of perfused vasculature with age. Wholemount imaging with light-sheet microscopy of the renal arterial vasculature perfused with Evans blue in 7-month-old control (*Ntn1^fl/fl^ or Foxd1^GC/+^;Ntn1^fl/+^*) and mutant (*Foxd1^GC/+^;Ntn1^fl/fl^*) mice (**A**). Quantification of wholemount imaging for vascular branching and total vascular area (µm x10^6^) in 2-month versus 7-month-old male mice (**B**). Light-sheet imaging of whole kidneys was performed using a LaVision UltraMicroscope II and 3D reconstruction the vascular network and quantification was conducted using Imaris imaging software. Two-way ANOVA, n=3-6 per group. Glomerular counts (**C**), plasma creatinine (mg/dL) and blood urea nitrogen (BUN, mg/dL) (**D**) in adult control and mutant mice, t-test, n=3 per group. Control mice are shown in white bars and mutant mice are shown in gray bars. Values are mean±SEM or SD. Statistical significance p<0.05 (*).

## Discussion

In this study we demonstrate that proper patterning of the renal interlobar arteries is not required for normal kidney function. Further, our results suggest that it is possible for arterial patterning to be more efficient and aid in improving response to ischemic injury or age-related vascular loss. These data expand on previous studies which identified Netrin-1, secreted by stromal progenitors during nephrogenesis, as an essential regulator of arterial patterning and maturation^8,9^. Importantly, the results from the current study highlight that our understanding of renal vasculature remains incomplete. Understanding how patterning and maturation of the arterial tree affects physiology and response to injury or aging is likely to have important implications for enhancing kidney regeneration and tissue engineering.

Netrin-1 is a known axonal guidance cue that has also been shown to play a role in angiogenesis^21–24^. In the developing kidney, netrin-1 is secreted by Foxd1+ stromal progenitors during kidney development^8,9^. The physiological response observed in the current study is independent of the loss of this source of netrin-1 as the Foxd1+ progenitors are depleted around post-natal day 4. Rather, the protective effect observed in our netrin-1 (*Ntn1*) mutant mice appears to result from the changes to patterning or maturation of the renal vasculature. In the uninjured kidney of adult mice, netrin-1 has low levels of secretion from tubular epithelial cells, however following kidney injury, netrin-1 secretion from the tubules is greatly increased^28–30^. In human kidney injury, urinary netrin-1 is similarly increased^31,32^. This endogenous increase in netrin-1 following injury is thought to play a protective role by modulating the immune response and promoting regeneration^29–31^. Importantly, this source of tubular netrin-1 remains intact in our mutant mice, as our deletion specifically targets only the interstitial progenitors during development. While the underlying molecular mechanisms mediating this protection remains unknown, future studies aim to identify whether this effect is a result of alterations in blood flow due to the differential patterning or a result of molecular changes to the vascular wall, i.e. endothelial cells or vascular smooth muscle cells.

Changes in blood flow during early reperfusion have previously been shown to play an important role in the severity of tubular injury in the kidneys following ischemia^3^. Honeycutt *et al* and Luo *et al* reported a delay in vascular smooth muscle cell (VSMC) coverage of the arterial tree in netrin-1 (*Ntn1*) mutant mice^8,9^. VSMCs are a critical component of the arterial wall, and their contractile properties allow for tight control of blood flow and blood pressure^33,34^. It remains unknown if there are changes to VSMC density along the renal arterial tree in adult mutant mice. However, changes in VSMC density or function following a delay in their coverage during nephrogenesis could result in alterations to management of vascular tone under physiology stress (i.e. ischemia or again). Changes in perfusion have also been suggested to play a role in age-related vascular loss in the kidney through arterial stiffening^35,36^. Endothelial dysfunction, oxidative stress, and inflammation are all thought to contribute to stiffening of the arterial tree with age^35,36^. However, given we did not see changes in circulating or kidney resident immune cells at baseline with aging, we speculate that a reduction in inflammation is unlikely to drive the protection we observed in mice with an abnormal vascular pattern. Whether the abnormal pattern of the arterial tree observed in our netrin-1 (*Ntn1*) mutant mice alters baseline or post-ischemic perfusion to reduce, or prevent, either of these mechanisms warrants further investigation.

Loss of vascular density, or microvascular rarefaction, is known to play a critical role in kidney function and progression to chronic kidney disease (CKD) following ischemic kidney injury^37^. In studies which utilize rodent models and mimic what is observed in the clinical setting, loss of cortical peritubular or outer-medullary plexus capillaries is often observed in association with severe tubular injury following periods of renal ischemia^37,38^. It is thought that capillary loss contributes to non-recovery and progression to CKD through impaired hemodynamics and prolonged inflammation, ultimately leading to glomerulosclerosis and tubulointerstitial fibrosis^37,39,40^. Similarly, loss of capillary density is often observed with aging^41–44^. In the kidney, vascular aging has also been shown to drive CKD progression^42,45^. The protection associated with the abnormal vascular pattern in mutant mice was observed in both sexes, despite females having less ischemic injury than males. The further decrease in tubular injury in female mutant mice have important implications for sex-specific mediators of CKD. This is of particular importance as studies have shown that more women than men are diagnosed with CKD, but female patients are protected from the development of end-stage kidney disease^46,47^. The data from the current study suggests that it is possible to prevent loss of vascular density and maintain vessel integrity. As such, the protection observed in our netrin-1 (*Ntn1*) male and female mutant mice may provide insight into the factors which positively regulate vascular density in order to prevent capillary drop-out.

While the current data shows a reduction in kidney injury in mutant mice with an abnormal vascular pattern, there is variability in the response. Our previous study also found variation in the arterial patterning between netrin-1 (*Ntn1*) mutant mice^8^. Whole-mount imaging of the vascular tree revealed the interlobar arteries of some mutant mice show only a slight variation from wild-type, while the patterning in others was completely stochastic. We speculate that this variation in renal vascular patterning contributes to the range of injury in mutant mice. Such that, the mutant mice with higher levels of injury likely have vascular patterning that is more similar to wild-type. Future studies utilizing Evans blue perfusion of the renal vasculature post-ischemia can be performed to investigate this association. Additionally, selection of a moderate (26-minute) period of ischemia may also contribute to the variability of injury in mutant mice as it is known that moderate levels of ischemia result in greater variability in injury than using more severe ischemic times^48–51^.

To date, our characterization of vascular patterning in netrin-1 (*Ntn1*) mutant mice has focused on the arterial tree and the effect on the renal veins is currently elusive. The patterning of the renal veins is of interest as they have been implicated in playing a significant role in ischemic injury^3,52^. However, to date, there are no appropriate markers to specifically label the venous vessels in the kidney. Whole-mount imaging to visualize the vasculature utilizing SMA requires significant VSMC coverage, and therefore was an unreliable marker for venous vessels. Co-labeling vessels with VSMCs and endothelial marker CD31, was also insufficient to label the large venules and veins in the kidney. Similarly, perfusion with Evans blue did not label the vasculature past the glomeruli. While this remains unknown, we speculate that the renal veins are patterned in parallel with the arterial tree, particularly the larger arcuate and interlobar vessels. Therefore, altered venous patterning in the netrin-1 (*Ntn1*) mutant mice could also contribute to the observed protective effect through improved blood flow or a reduction in venous pressures to the kidney.

To our knowledge, this is the first study to investigate the impact of renal vascular patterning on kidney function. While it was surprising that netrin-1 (*Ntn1*) mutant mice with a stochastic arterial tree were protected from ischemic injury and age-related vessel loss, we are now able to utilize this model to investigate mechanisms of vascular integrity. The current study also has important implications for vascularization of kidney organoids and *de novo* kidney tissue engineering, which have proved challenging to form appropriate vessel networks. These data suggest that “normal” patterning of these vessels may not be necessary in order to for appropriate function. In fact, it may be possible to guide the vasculature to a more efficient pattern. Understanding how vascular patterning affects a number of physiological responses in the kidney provides critical insight for identifying novel therapeutic targets for clinically relevant approaches to treat kidney disease.

## Methods

### Animals

All animal studies were approved by the Office of Animal Care and Use at the University of North Carolina at Chapel Hill (UNC-CH) and the UNC-CH Institutional Animal Care and Use Committee (IACUC). Animal husbandry is performed by UNC’s Division of Comparative Medicine (DCM). All animals for this study were kept in a temperature and humidity-controlled environment with a 12-hour light/dark cycle. No more than five adult animals were housed per cage. All mice had free access to food (LabDiet PicoLab Select Rodent) and water. All mouse lines were maintained on the C57Bl6/J background (Jackson Labs, Strain #000664). Details of genetically edited mouse lines utilized are as follows: *Foxd1^GC/+^*(Cre Strain #012463)^8,53^ and *Ntn1^fl/fl^* (Floxed Strain #028038)^8,54^ strains were utilized. *Foxd1^GC/+^;Ntn1^fl/fl^* mutant mice were generated by crossing a Cre-expressing *Foxd1* male mouse to a *Ntn1* floxed female mouse. Wild-type (WT) mice were either *Ntn1^fl/fl^* or *Ntn1^fl/^.* Heterozygous (Het) mice were *Foxd1^GC/+^;Ntn1^fl/+^*. For IR experiments, WT and Het mice were combined into a single control group as we did not find differences between genotypes for any of the analyses. Genotyping primers are listed in **Table S1**. Male and female mice were run in parallel to appropriately analyze the data for sex differences.

### Ischemia-Reperfusion surgery

For each surgery, all surgical instruments were sterilized by autoclave. Mice were anesthetized with inhaled isoflurane (∼4% induction, ∼2% maintenance). Ophthalmic lubricant was applied to the eyes to prevent dryness. Following confirmation anesthesia (toe pinch), the hair was shaved from the abdomen and the surgical site was cleaned with alternating betadine and 70% ethanol (3 times). The mice were then placed on a servo-controlled heating table to maintain a temperature at 37°C. The kidneys were approached via a small dorsal flank incision. The upper and lower poles of both kidneys and the renal pedicle were carefully dissected from surrounding tissue by blunt dissection with forceps. An atraumatic microvascular clamp (Fine Science Tools 18052-03) was then applied to each renal pedicle (artery and vein), occluding blood flow to/from the kidneys for 26-minutes. A clamp was first applied to one renal pedicle, and as quickly as possible (within approximately 2 mins) a clamp was applied to the other renal pedicle to ensure bilateral ischemia. The body cavity and skin incision were covered during the ischemic time with sterile saline-soaked gauze. Following ischemia, the clamps were removed, and restoration of blood flow to the kidneys was confirmed visually by a change in color from dusky purple to pink. Sterile saline (approximately 0.2 ml) was introduced into the retroperitoneal space to help replace any fluids lost and the wound closed (5-0 Vicryl absorbable suture for the muscle layer, 6-0 Monofilament Nylon Nonabsorbable Suture for the skin). Bupivacaine and Meloxicam 5mg/kg (diluted with sterile saline to 0.5mg/ml then adjusted to body weight) were administered for analgesia. As the anesthesia was withdrawn, the mice were monitored until fully conscious and allowed to recover in a warm, clean cage with free access to food and water. The mice were humanely euthanized following one-day of recovery.

### Metabolic cages

Mice were housed in metabolic cages for 24-hour urine collection following an overnight acclimation period. Metabolic cage experiments were performed in an AAALAC approved UNC animal facility in a temperature and humidity-controlled environment. Normal 12-hour light/dark cycles were maintained. Mice had free access to food and water for the duration of the study. 24-hour urine was collected, centrifuged (4000 RPM, 10-minutes), and stored at -80°C until analyses were performed.

### Tissue Harvest

Mice were deeply anesthetized with isoflurane (approximately 4-5%), and a midline abdominal incision performed. Blood was collected from the abdominal aorta into tubes coated with 100 mM EDTA (pH 7.4). The kidneys were then rapidly excised. The right kidney was snap frozen in liquid nitrogen and stored at -80°C. The left kidney was cut at midline and one-half was fixed in 10% neutral buffered formalin (VIP-Fixative, Fisher 23–730–587) for histologic analysis. The other one-half of the left kidney was fixed in 4% paraformaldehyde for 1 hour, washed with 1X PBS (3 times), maintained overnight in 30% sucrose and embedded in OCT compound for sectioning. Blood was centrifuged (3500 RPM, 10-minutes) and plasma stored at -80°C. Ear clip to confirm genotyping was performed prior to weaning and at euthanasia for each animal.

### Assessment of Kidney Function

Creatinine (Cr) concentration in urine and plasma and blood urea nitrogen (BUN) concentrations were determined by a QuantiChrom Cr assay kit (BioAssay Systems, DICT 500) and QuantiChrom urea assay kit (BioAssay Systems, DIUR 100), respectively, according to the manufacturer’s instructions. Proteinuria was measured in 24-hour urine using a Bradford Assay. Creatinine clearance was determined using the following equation: Creatinine Clearence (GFR) = (Urine**_Cr_** x 24h Urine Volume) / Plasma**_Cr_**

### Periodic Acid Schiff

Periodic Acid Schiff (PAS, Abcam ab150680) staining was performed on formalin-fixed (VIP-Fixative, Fisher 23-730-587), paraffin-embedded kidney sections (3µm) per the manufacturer’s instructions. Briefly, slides with kidney sections were deparaffinized in xylene and rehydrated with graded ethanol and water. Slides were then placed in Periodic Acid Solution for 10-minutes and rinsed with deionized water (4 times). Slides with kidney sections were then immersed in Schiff’s solution for 30-minutes before washing in warm, running tap water for 10-minutes. Slides were next immersed in Hematoxylin for 3-minutes, rinsed briefly with water, and incubated in Bluing reagent for 30-seconds. Slides with kidney sections were placed in a final rinse with water, dehydrated with 100% ethanol, re-immersed in xylene, and mounted (Cytoseal XYL, Fisher 22-050-262).

### Tubular Injury Scoring

PAS-stained slides were scored by research collaborators who were expertly trained in kidney injury pathology as previously reported^26^ and who were blinded to both sample identifiers and study hypotheses. Large cortical and outer medullary stitched images were captured with Axio Imager by tiling nine 20x images (9 images: 3x3) per slide. Each image was divided into 144 areas and then assessed if the injured tubules are >50% of one small area (these areas are considered as positive injured area). Renal damage including dead tubules, loss of brush borders, tubule dilatation, cast formation, tubular epithelial swelling, and vacuolar degeneration was semi quantitatively scored. Score 0 represents injury area ≤ 5%, scores 1, 2, 3, 4, and 5 exhibit the damage involving 5% to 30%; 31% to 50%; 51% to 70%; 71% to 90% and >90% of the whole kidney area, respectively. Representative histology images were taken with a Leica DMi8 inverted widefield fluorescent microscope with Leica DFC9000 GT camera and LASX software.

### Immunofluorescent staining and Imaging

Slides with kidney sections (8-10µm) for immunofluorescent staining were removed from the −80°C freezer and equilibrated to room temperature. Sections were rehydrated, and OCT was removed by incubating the slides in 1x PBS for 10-minutes. A hydrophobic barrier was drawn around the section using a histology pen (Vector Laboratories H-4000) and sections were incubated for 45-minutes in blocking solution (1X PBS, 4% donkey serum (Equitech-Bio SD30-0100), 1% bovine serum albumin (Fisher BP9706-100), and 0.25% Triton X-100 (Fisher BP151-500)). Sections were incubated at room temperature for 2 hours with primary antibodies (Megalin/LRP2, MyBiosource MBS690201, 1:500) (KIM-1, R&D Systems MAB1817, 1:250) in block solution. Slides were washed (5-minutes, 3x) at room temperature wash solution (1X PBS + 0.25% Triton X-100). Sections were then stained with Alexa Fluor secondary antibodies (488, 568, or 647 conjugated) in blocking solution for 1-hour. Finally, slides were washed (5-minutes, 3x) covered from light using wash solution with the last wash containing DAPI diluted at 1:10000. Slides were mounted using Prolong gold mounting media (ThermoFisher P36930). Images were captured with a Leica DMi8 inverted widefield fluorescent microscope with Leica DFC9000 GT camera and LASX software. Images were taken at 5x magnification and acquisition settings were the same for all samples.

### Evans blue staining and whole mount imaging

Evans blue staining and imagine was performed as previously described^8,27^. Briefly, mice were anesthetized under isoflurane (∼2%) and surgical plane was confirmed by toe pinch. The chest cavity and peritoneum are cut to expose the diaphragm. Evans blue dye (2%) in 0.9% saline at 3 μl/g of body weight was injected into the left ventricle through the diaphragm. The dye circulated (∼2-5 minutes) until the exposed areas (paws, tail, chin) turn blue. The animal is subsequently euthanized. Evans blue-stained kidneys were then collected, rinsed briefly in 1X PBS several times to remove excess dye, and fixed in 4% PFA overnight at 4°C. Samples are washed in 1X PBS (3 times) and stored at 4°C until clearing.

### Tissue clearing and imaging

Tissue clearing of Evans blue-stained kidneys was performed via a modified iDISCO protocol^55^. All following steps are performed with rocking unless otherwise noted. Adult kidneys were dehydrated in increasing methanol (MeOH)/H2O series of 25%, 50%, 75%, and 100% for 1 hour at RT. Samples were then incubated in 66% dichloromethane (DCM)/33% MeOH for 3 hours at RT. Samples were then processed through two rounds of incubation with 100% DCM at RT. DCM is removed, and 100% dibenzyl ether (DBE) is added. Samples are incubated, with no rocking, until clear (+ 24 hours). Samples are stored in 100% DBE until imaging. Light-sheet imaging of whole kidneys was performed using a LaVision UltraMicroscope II (LaVision BioTec). 3D reconstruction of images and vascular network quantification was conducted using Imaris imaging software. Analysis of the 2-month vascular metrics was previously reported^8^. These data have been re-analyzed in Figure 5 for age-related comparisons to 7-month-old mice.

### Quantification of wholemount 3D vasculature

Quantification of renal vasculature was performed with Imaris as previously described^8^. Briefly, a surface of the 3D image was first created to visualize the stained vasculature. A binary mask of the rendered surface was created using the “Mask channel” function. “Constant inside/outside” settings for voxel intensity was set to 0 outside and 100 inside which was necessary to render the vasculature solid in order for the filament tracer module within Imaris to develop wireframes for statistical analysis. Supervised automatic tracing was performed to ensure that automatically generated paths were not duplicated, or artifacts inaccurately rendered as vasculature. As both kidneys from the same animal had similar statistical values with little deviation when averaged, only one kidney per animal was quantified. This method allowed unbiased quantification of the renal arterial trees for Evans blue-stained kidneys from adult mice. For graphical display of vascular patterning and imaging that were not quantified, a surface was created for the desired channel. This surface was then masked using the same procedure, but “Constant inside/outside” settings for voxel intensity were set to 0 outside with the inside setting left unchecked.

### Glomerular Counts

Glomerular counts were performed following the acid maceration protocol from Peterson *et al*^56^. In brief, kidneys were collected and weighed, then bifurcated lengthwise. Each kidney half was then cut into roughly 2mm pieces and transferred to a 15ml conical tube. 5ml of 6M HCl acid was added to each tube. Tubes were then gently agitated and incubated in 37°C water bath for 90 minutes (gentle agitation every 15 minutes). Following incubation, samples were homogenized by backfilling a 5ml syringe with the kidney-acid solution then extruding through an 18G needle and repeated through a 21G needle. PBS was then added to the kidney-acid solution to bring the total volume to 50ml. Samples were incubated overnight on a rocker at 4°C and glomeruli counted within 2 days of processing. To count glomeruli, the tube was inverted to mix the sample, 500µl of the kidney solution and 500µl of PBS were added to a 12 well plate. The total number of glomeruli per well was counted using a Leica DMi8 inverted widefield microscope with a Leica DFC9000 GT camera and LASX software. The glomeruli were counted in triplicate then averaged. Counting was blinded and performed twice by independent researchers.

### Whole Blood Analysis

250ul of whole blood was collected from each animal by submandibular puncture at the required timepoints in EDTA coated tubes. The whole blood was immediately analyzed via an IDEXX ProCyte Dx veterinary CBC hematology analyzer.

### Statistics

Data were analyzed using GraphPad Prism software (GraphPad Inc.). Baseline and post-ischemia data were analyzed by two-way ANOVA to compare genotype and sex. Data in aged mice were analyzed using two-way ANOVA or Student’s unpaired *t* test. Specific statistical analysis utilized for each dataset is listed in figure legends. Significance was determined by *P*<0.05.

## Acknowledgements

The authors would like to thank the members of the O’Brien lab for technical help and critical discussions, Dr. Pablo Ariel, director of the UNC Microscopy Services Laboratory (MSL), for assistance with light-sheet imaging and Imaris rendering/analysis, Dr. Arendshorst (retired), Dr. Taylor, and Xi Yang for sharing equipment for surgical procedures and metabolic cages, the histology research core and pathology core for processing and sectioning of kidney tissues, and our funding, NIDDK R01DK121014 to LO (PI), NIDDK R01DK133369 to TS (PI) and LO (Co-I), and NIDDK U2CDK133491 (trainee funding for SRM). The MSL is supported by the P30CA016086 Cancer Center Core Support Grant to the UNC Lineberger Comprehensive Cancer Center. Light-sheet microscopy at MSL is also supported, in part, by the North Carolina Biotech Center Institutional Support Grant 2016-IDG-1016.

## Supplemental Figure Legends

**Supplemental Table 1.**
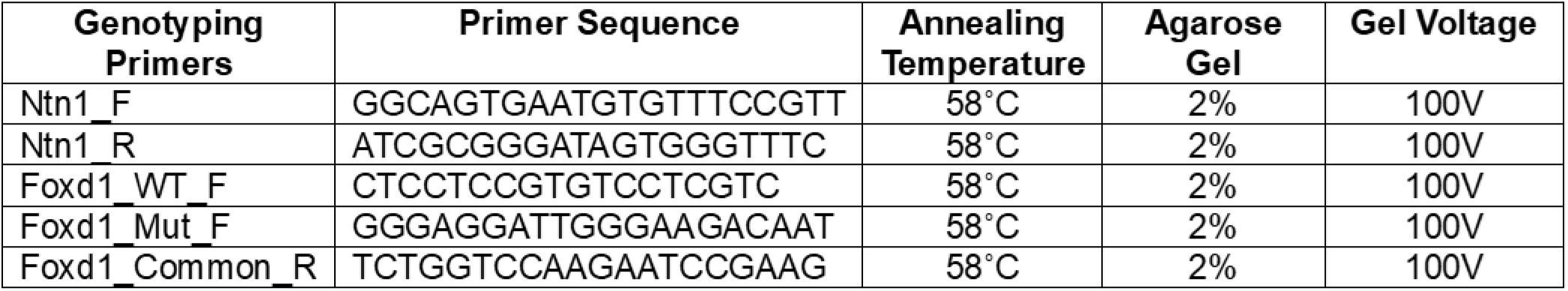
Primer sequences for genotyping. Genotyping primer name (column 1), primer sequences (column 2), annealing temperatures (column 3), percent (%) agarose gel (column 4), and gel running voltage (column 5). Abbreviations: Netrin 1 floxed allele (Ntn1), Foxd1 Cre (Foxd1), forward primer (F), reverse primer (R). All primers sequences were obtained from Jax Laboratories and confirmed with positive and negative controls. Mice are genotyped prior to weaning and at euthanasia.

**Supplemental Figure 1.**
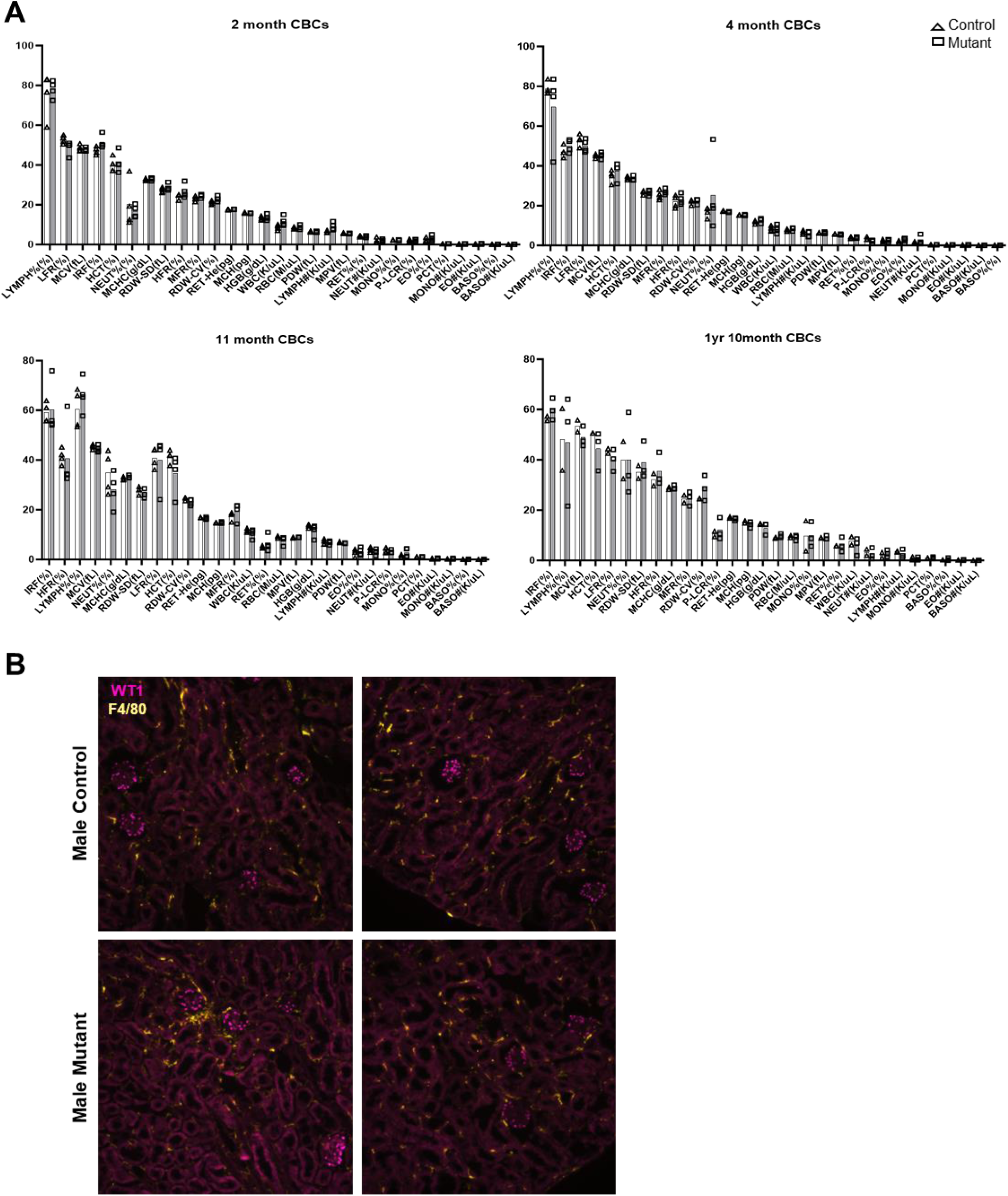
Circulating and kidney immune cells. Complete blood count based inflammatory score (CBCs) in control (*Ntn1^fl/fl^ or Foxd1^GC/+^;Ntn1^fl/+^*) and mutant (*Foxd1^GC/+^;Ntn1^fl/fl^*) mice at 2, 4, 11, and 22 months of age (**A**). Control mice are shown in white bars and mutant mice are shown in gray bars. Values for each animal is shown and are expressed at percent (%) or number (#) of cells or concentration (pg, g/dL, K/µl, fL). N= 2-4 mice per group. Abbreviations: lymphocytes (LYMPH), reticulocyte low fluorescent ratio (LFR), mean corpuscular volume (MCV), immature reticulocyte fraction (IRF), hematocrit (HCT), neutrophils (NEUT), mean corpuscular hemoglobin concentration (MCHC), red blood cell distribution width (RDW-SD/CV), reticulocyte high fluorescent ratio (HFR), reticulocyte middle fluorescent ratio (MFR), reticulocyte hemoglobin content (RET-He), mean corpuscular hemoglobin (MCH), hemoglobin (HGB), white blood cells (WBC), red blood cells (RBC), (PDW), mean platelet volume (MPV), reticulocytes (RET), monocytes (MONO), platelet-large cell ratio (p-LCR), eosinophils (EO), plateletcrit (PCT), basophils (BASO). Representative immunofluorescent images of F4/80 monocytes/macrophages in the kidney from control (*Ntn1^fl/fl^ or Foxd1^GC/+^;Ntn1^fl/+^*) and mutant (*Foxd1^GC/+^;Ntn1^fl/fl^*) mice at baseline (∼4 months old) (**B**). Wilms tumor (WT1) positive podocytes (magenta) and F4/80 positive monocytes/macrophages (yellow). Representative images are from the renal cortex of two mice. N=3-8 mice per group, kidneys are from mice used for injury analysis in figure 1. Images were captured with a Leica DMi8 inverted widefield fluorescent microscope with Leica DFC9000 GT camera and LASX software. Images were taken at 20x magnification and acquisition settings were the same for all samples.

**Supplemental Figure 2.**
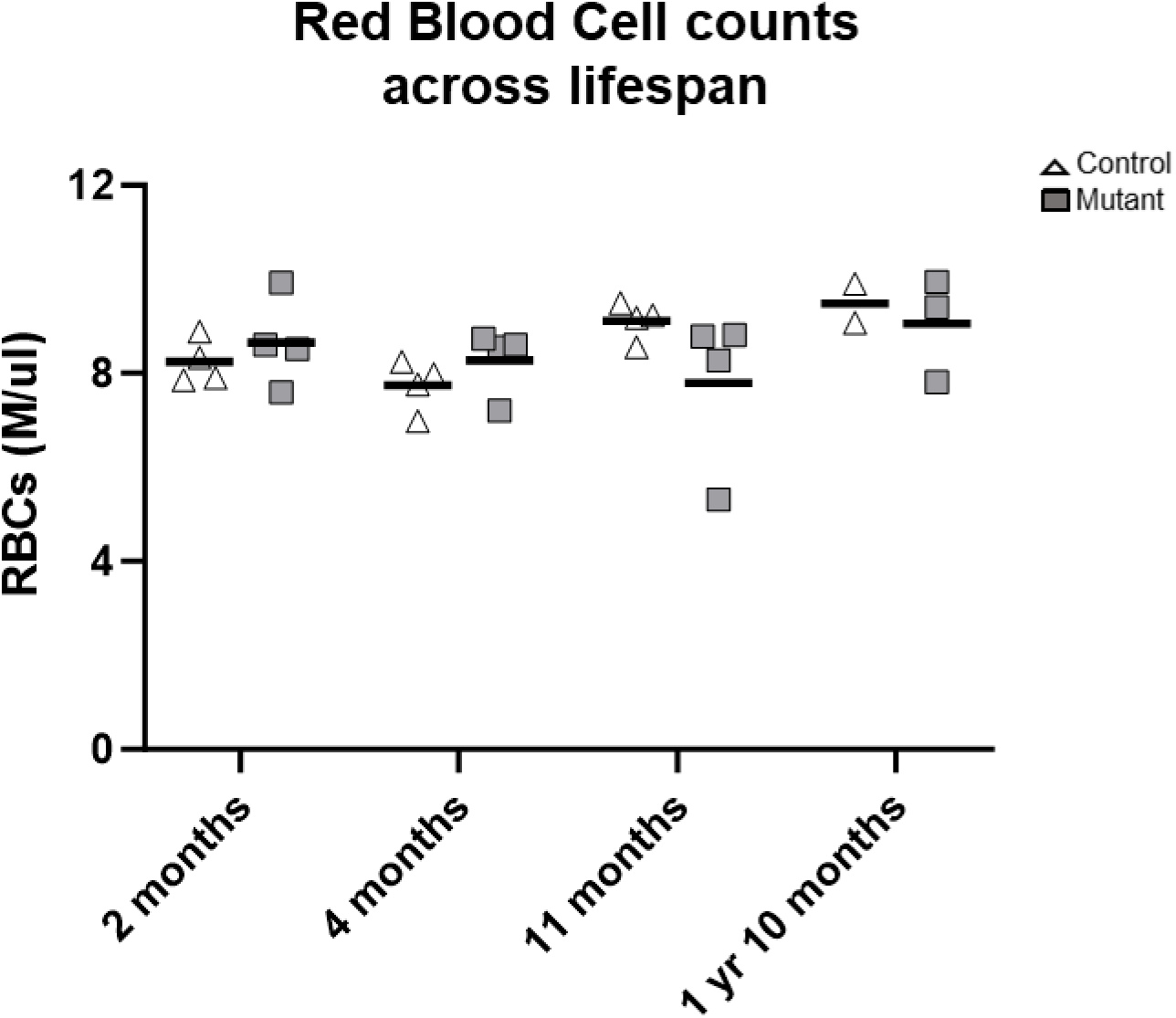
Red blood cell counts. Red blood cell (RBC) counts in control (*Ntn1^fl/fl^ or Foxd1^GC/+^;Ntn1^fl/+^*) and mutant (*Foxd1^GC/+^;Ntn1^fl/fl^*) mice at 2, 4, 11, and 22 months of age. Control mice are shown in white bars and mutant mice are shown in gray bars. Values for each animal is shown, horizontal line indicates mean. Concentration is shown as millions of cells per microliter (M/µl). N= 2-4 mice per group and are the same animals from the CBC panel in supplemental figure 1.

**Supplemental Figure 3.**
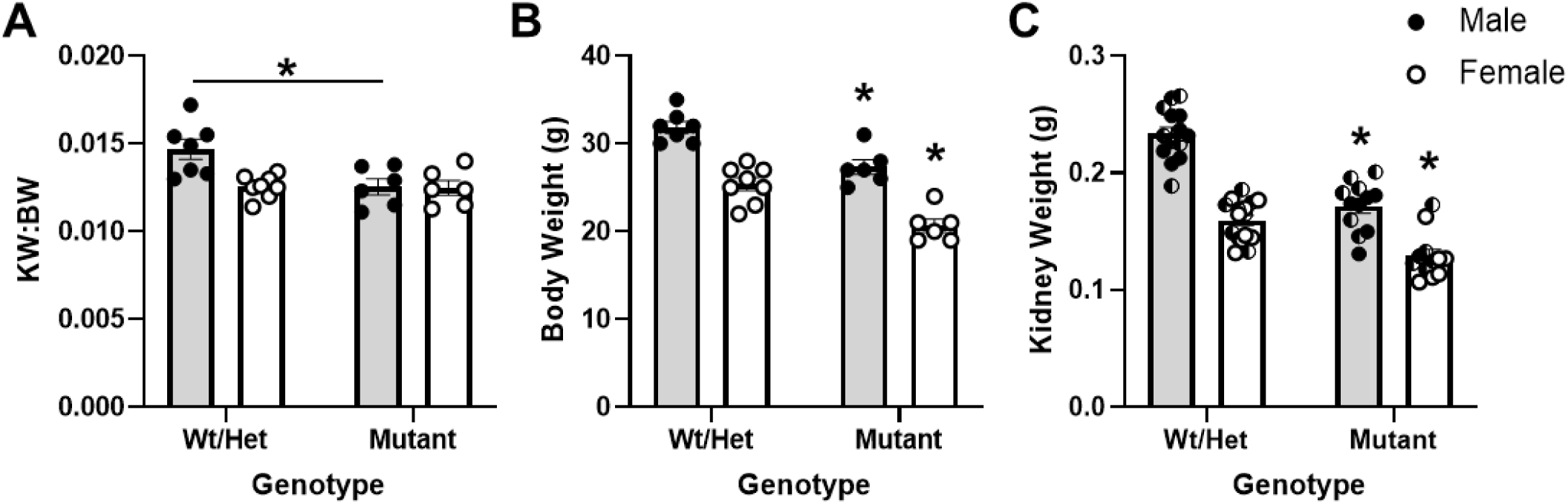
Body and kidney weight metrics post-ischemia. Kidney to body weight ratio (KW:BW) (**A**), body weight (g) (**B**), and kidney weight (g) (**C**) following a 26-minute bilateral ischemia and one-day of recovery in wild-type (Wt, *Ntn1^fl/fl^*) and heterozygous (Het, *Foxd1^GC/+^;Ntn1^fl/+^*) compared to mutant (*Foxd1^GC/+^;Ntn1^fl/fl^*) adult mice (∼4 months old). Wt and Het mice were combined into a control group as no differences were observed. Male mice are represented in gray bars and female mice are represented in white bars. Values are mean±SEM, n=6-8 per group. Weights are from mice used for injury analysis in figure 3. Two-way ANOVA, statistical significance p<0.05 (*).

## References

1. Kriz, W. Structural organization of renal medullary circulation. Nephron 31, 290–295 (1982).

2. Evans, R. G., Eppel, G. A., Anderson, W. P. & Denton, K. M. Mechanisms underlying the differential control of blood flow in the renal medulla and cortex. J. Hypertens. 22, 1439 (2004).

3. McLarnon, S. R. et al. Lipopolysaccharide Pretreatment Prevents Medullary Vascular Congestion following Renal Ischemia by Limiting Early Reperfusion of the Medullary Circulation. J. Am. Soc. Nephrol. 33, 769 (2022).

4. Molema, G. & Aird, W. C. Vascular Heterogeneity in the Kidney. Semin. Nephrol. 32, 145–155 (2012).

5. Munro, D. A. D., Hohenstein, P. & Davies, J. A. Cycles of vascular plexus formation within the nephrogenic zone of the developing mouse kidney. Sci. Rep. 7, 3273 (2017).

6. Daniel, E. et al. Spatiotemporal heterogeneity and patterning of developing renal blood vessels. Angiogenesis 21, 617–634 (2018).

7. Dumas, S. J. et al. Phenotypic diversity and metabolic specialization of renal endothelial cells. Nat. Rev. Nephrol. 17, 441–464 (2021).

8. Honeycutt, S. E. et al. Netrin 1 directs vascular patterning and maturity in the developing kidney. Dev. Camb. Engl. 150, dev201886 (2023).

9. Luo, P. M., Gu, X., Chaney, C., Carroll, T. & Cleaver, O. Stromal netrin 1 coordinates renal arteriogenesis and mural cell differentiation. Development 150, dev201884 (2023).

10. McMahon, A. P. Development of the Mammalian Kidney. Curr. Top. Dev. Biol. 117, 31–64 (2016).

11. Long, D. A., Norman, J. T. & Fine, L. G. Restoring the renal microvasculature to treat chronic kidney disease. Nat. Rev. Nephrol. 8, 244–250 (2012).

12. Nangaku, M. Chronic hypoxia and tubulointerstitial injury: a final common pathway to end-stage renal failure. J. Am. Soc. Nephrol. JASN 17, 17–25 (2006).

13. Tanaka, S., Tanaka, T. & Nangaku, M. Hypoxia as a key player in the AKI-to-CKD transition. Am. J. Physiol. Renal Physiol. 307, F1187–1195 (2014).

14. Afsar, B. et al. Capillary rarefaction from the kidney point of view. Clin. Kidney J. 11, 295–301 (2018).

15. Steegh, F. M. E. G. et al. Capillary rarefaction: a missing link in renal and cardiovascular disease? Angiogenesis 27, 23–35 (2024).

16. Lindenmeyer, M. T. et al. Interstitial vascular rarefaction and reduced VEGF-A expression in human diabetic nephropathy. J. Am. Soc. Nephrol. JASN 18, 1765–1776 (2007).

17. Qian, Q. Inflammation: A Key Contributor to the Genesis and Progression of Chronic Kidney Disease. Contrib. Nephrol. 191, 72–83 (2017).

18. Lu, J. et al. Discrete functions of M2a and M2c macrophage subsets determine their relative efficacy in treating chronic kidney disease. Kidney Int. 84, 745–755 (2013).

19. Gewin, L., Zent, R. & Pozzi, A. Progression of chronic kidney disease: too much cellular talk causes damage. Kidney Int. 91, 552–560 (2017).

20. Geuens, T., van Blitterswijk, C. A. & LaPointe, V. L. S. Overcoming kidney organoid challenges for regenerative medicine. NPJ Regen. Med. 5, 8 (2020).

21. Serafini, T. et al. The netrins define a family of axon outgrowth-promoting proteins homologous to C. elegans UNC-6. Cell 78, 409–424 (1994).

22. Kennedy, T. E., Serafini, T., de la Torre, J. R. & Tessier-Lavigne, M. Netrins are diffusible chemotropic factors for commissural axons in the embryonic spinal cord. Cell 78, 425–435 (1994).

23. Larrivée, B. et al. Activation of the UNC5B receptor by Netrin-1 inhibits sprouting angiogenesis. Genes Dev. 21, 2433–2447 (2007).

24. Tu, T. et al. CD146 acts as a novel receptor for netrin-1 in promoting angiogenesis and vascular development. Cell Res. 25, 275–287 (2015).

25. Lu, X. et al. The netrin receptor UNC5B mediates guidance events controlling morphogenesis of the vascular system. Nature 432, 179–186 (2004).

26. Ide, S. et al. Ferroptotic stress promotes the accumulation of pro-inflammatory proximal tubular cells in maladaptive renal repair. eLife 10, e68603 (2021).

27. Honeycutt, S. E. & O’Brien, L. L. Injection of Evans blue dye to fluorescently label and image intact vasculature. BioTechniques 70, 181–185 (2021).

28. Brian Reeves, W., Kwon, O. & Ramesh, G. Netrin-1 and kidney injury. II. Netrin-1 is an early biomarker of acute kidney injury. Am. J. Physiol.-Ren. Physiol. 294, F731–F738 (2008).

29. Ranganathan, P., Jayakumar, C. & Ramesh, G. Proximal tubule-specific overexpression of netrin-1 suppresses acute kidney injury-induced interstitial fibrosis and glomerulosclerosis through suppression of IL-6/STAT3 signaling. Am. J. Physiol.-Ren. Physiol. 304, F1054–F1065 (2013).

30. Mohamed, R., Jayakumar, C., Ranganathan, P. V., Ganapathy, V. & Ramesh, G. Kidney Proximal Tubular Epithelial-Specific Overexpression of Netrin-1 Suppresses Inflammation and Albuminuria through Suppression of COX-2-Mediated PGE2 Production in Streptozotocin-Induced Diabetic Mice. Am. J. Pathol. 181, 1991–2002 (2012).

31. Reeves, W. B., Kwon, O. & Ramesh, G. Netrin-1 and kidney injury. II. Netrin-1 is an early biomarker of acute kidney injury. Am. J. Physiol. Renal Physiol. 294, F731-738 (2008).

32. Ramesh, G., Kwon, O. & Ahn, K. Netrin-1: a novel universal biomarker of human kidney injury. Transplant. Proc. 42, 1519–1522 (2010).

33. Shi, J., Yang, Y., Cheng, A., Xu, G. & He, F. Metabolism of vascular smooth muscle cells in vascular diseases. Am. J. Physiol.-Heart Circ. Physiol. 319, H613–H631 (2020).

34. Touyz, R. M. et al. Vascular smooth muscle contraction in hypertension. Cardiovasc. Res. 114, 529–539 (2018).

35. Tanriover, C. et al. Early aging and premature vascular aging in chronic kidney disease. Clin. Kidney J. 16, 1751–1765 (2023).

36. Hobson, S. et al. Accelerated Vascular Aging in Chronic Kidney Disease: The Potential for Novel Therapies. Circ. Res. 132, 950–969 (2023).

37. Basile, D. P., Anderson, M. D. & Sutton, T. A. Pathophysiology of acute kidney injury. Compr. Physiol. 2, 1303–1353 (2012).

38. Eardley, K. S. et al. The role of capillary density, macrophage infiltration and interstitial scarring in the pathogenesis of human chronic kidney disease. Kidney Int. 74, 495–504 (2008).

39. Menshikh, A. et al. Capillary rarefaction is more closely associated with CKD progression after cisplatin, rhabdomyolysis, and ischemia-reperfusion-induced AKI than renal fibrosis. Am. J. Physiol. - Ren. Physiol. 317, F1383–F1397 (2019).

40. Scarfe, L. et al. Long-term outcomes in mouse models of ischemia-reperfusion-induced acute kidney injury. Am. J. Physiol. - Ren. Physiol. 317, F1068–F1080 (2019).

41. Chade, A. R. Renal Vascular Structure and Rarefaction. Compr. Physiol. 3, 817–831 (2013).

42. Fang, Y. et al. The ageing kidney: Molecular mechanisms and clinical implications. Ageing Res. Rev. 63, 101151 (2020).

43. O’Sullivan, E. D., Hughes, J. & Ferenbach, D. A. Renal Aging: Causes and Consequences. J. Am. Soc. Nephrol. JASN 28, 407–420 (2017).

44. Faber, J. E. et al. Aging Causes Collateral Rarefaction and Increased Severity of Ischemic Injury in Multiple Tissues. Arterioscler. Thromb. Vasc. Biol. 31, 1748–1756 (2011).

45. Zhang, Y., Yu, C. & Li, X. Kidney Aging and Chronic Kidney Disease. Int. J. Mol. Sci. 25, 6585 (2024).

46. Lewandowski, M. J. et al. Chronic kidney disease is more prevalent among women but more men than women are under nephrological care. Wien. Klin. Wochenschr. 135, 89–96 (2023).

47. Carrero, J. J., Hecking, M., Chesnaye, N. C. & Jager, K. J. Sex and gender disparities in the epidemiology and outcomes of chronic kidney disease. Nat. Rev. Nephrol. 14, 151–164 (2018).

48. Dong, Y. et al. Ischemic Duration and Frequency Determines AKI-to-CKD Progression Monitored by Dynamic Changes of Tubular Biomarkers in IRI Mice. Front. Physiol. 10, 153 (2019).

49. Chen, Y.-L., Li, H.-K., Wang, L., Chen, J.-W. & Ma, X. No safe renal warm ischemia time—The molecular network characteristics and pathological features of mild to severe ischemia reperfusion kidney injury. Front. Mol. Biosci. 9, 1006917 (2022).

50. Crislip, G. R. et al. Ultrasound measurement of change in kidney volume is a sensitive indicator of severity of renal parenchymal injury. Am. J. Physiol. Renal Physiol. 319, F447–F457 (2020).

51. Skrypnyk, N. I., Harris, R. C. & de Caestecker, M. P. Ischemia-reperfusion Model of Acute Kidney Injury and Post Injury Fibrosis in Mice. J. Vis. Exp. JoVE 50495 (2013) doi:10.3791/50495.

52. Wei, J. et al. Role of intratubular pressure during the ischemic phase in acute kidney injury. Am. J. Physiol. - Ren. Physiol. 312, F1158–F1165 (2017).

53. Kobayashi, A. et al. Identification of a multipotent self-renewing stromal progenitor population during mammalian kidney organogenesis. Stem Cell Rep. 3, 650–662 (2014).

54. Bin, J. M. et al. Complete Loss of Netrin-1 Results in Embryonic Lethality and Severe Axon Guidance Defects without Increased Neural Cell Death. Cell Rep. 12, 1099–1106 (2015).

55. Renier, N. et al. iDISCO: a simple, rapid method to immunolabel large tissue samples for volume imaging. Cell 159, 896–910 (2014).

56. Peterson, S. M. et al. Estimation of Nephron Number in Whole Kidney using the Acid Maceration Method. J. Vis. Exp. JoVE (2019) doi:10.3791/58599.

